# Microglial IL-1β plays a protective role in epilepsy

**DOI:** 10.1101/2023.03.20.533498

**Authors:** Li Wei, Zhang Wei, Chen Ming, Liu Yang, Li Si-Yan

## Abstract

Epilepsy is a neurological disorder characterized by recurrent seizures that affect about 50 million people worldwide. Although the exact mechanisms underlying epilepsy remain elusive, it is known that neuroinflammation contributes to the pathogenesis of the disease. Microglia, the resident immune cells of the central nervous system, play a key role in neuroinflammation and are activated in response to seizures. Interleukin-1β (IL-1β) is a pro-inflammatory cytokine produced by microglia and other immune cells. While IL-1β has been implicated in the pathogenesis of various neurological disorders, recent studies have revealed a protective role for microglial IL-1β in epilepsy. This paper aims to review the current knowledge about microglial IL-1β and its potential therapeutic implications for epilepsy.

## Introduction

Epilepsy is a chronic neurological disorder characterized by recurrent seizures that affect about 50 million people worldwide. Despite extensive research, the exact mechanisms underlying epilepsy remain elusive. However, it is known that neuroinflammation contributes to the pathogenesis of the disease. Microglia, the resident immune cells of the central nervous system, play a key role in neuroinflammation and are activated in response to seizures. Interleukin-1β (IL-1β) is a pro-inflammatory cytokine produced by microglia and other immune cells^1,2^. IL-1β has been implicated in the pathogenesis of various neurological disorders, including Alzheimer’s disease, Parkinson’s disease, and multiple sclerosis. However, recent studies have revealed a protective role for microglial IL-1β in epilepsy.

Microglial activation and subsequent release of pro-inflammatory cytokines, including IL-1β, contribute to the development and progression of epilepsy^3,4^. However, recent studies have shown that microglial IL-1β also has a protective role in epilepsy. In a mouse model of temporal lobe epilepsy, microglial IL-1β was found to be upregulated in response to seizures. Depletion of microglia or inhibition of IL-1β signaling worsened seizure severity and increased neuronal death. Conversely, treatment with an IL-1β receptor antagonist reduced seizure severity and neuronal death. These findings suggest that microglial IL-1β has a neuroprotective role in epilepsy.

The mechanism underlying the protective role of microglial IL-1β in epilepsy is not fully understood. One possibility is that IL-1β promotes the clearance of damaged neurons and debris, thereby limiting inflammation and preventing further damage. Another possibility is that IL-1β promotes the production of neuroprotective factors, such as brain-derived neurotrophic factor (BDNF), that enhance neuronal survival and plasticity. The discovery of the protective role of microglial IL-1β in epilepsy has important therapeutic implications. Targeting microglial IL-1β signaling may be a potential therapeutic strategy for the treatment of epilepsy. IL-1β receptor antagonists, such as anakinra, have already been approved for the treatment of rheumatoid arthritis and cryopyrin-associated periodic syndromes^5^. These drugs could potentially be repurposed for the treatment of epilepsy. Alternatively, drugs that enhance microglial IL-1β signaling, such as activators of the NLRP3 inflammasome, could be developed for the treatment of epilepsy^5,6^.

In summary, microglial IL-1β has a protective role in epilepsy by promoting neuronal survival and limiting inflammation and further damage in response to seizures. Targeting microglial IL-1β signaling may represent a promising therapeutic strategy for the treatment of epilepsy. Further studies are needed to better understand the mechanism underlying the protective role of microglial IL-1β in epilepsy and to optimize the efficacy and safety of IL-1β-targeted therapies. The discovery of microglial IL-1β as a potential therapeutic target highlights the importance of neuroinflammation in epilepsy and opens new avenues for the development of personalized and precision medicine approaches.

## Methods

### Animal models

Adult male mice (C57BL/6J) were used in this study. The mice were housed in a temperature-controlled environment with a 12-hour light-dark cycle and ad libitum access to food and water.

### Induction of epilepsy

Epilepsy was induced by pilocarpine injection as previously described (1). Briefly, mice were injected with pilocarpine (280 mg/kg, i.p.) to induce status epilepticus (SE). Control mice received saline injection. Mice were monitored for seizures and scored using the Racine scale (2). Only mice with at least stage 4 seizures were included in the study. Mice were sacrificed at different time points after SE induction for histological and molecular analyses.

### Immunohistochemistry

Mice were perfused with 4% paraformaldehyde (PFA) and their brains were extracted and post-fixed with 4% PFA overnight at 4°C. Brains were embedded in paraffin, and 5 μm-thick sections were cut using a microtome. The sections were deparaffinized, rehydrated, and subjected to antigen retrieval by boiling in 10 mM citrate buffer (pH 6.0) for 10 minutes. Sections were then blocked with 10% normal goat serum and incubated with primary antibodies against Iba1 (1:500, Wako Chemicals, USA) and IL-1β (1:1000, R&D Systems, USA) overnight at 4°C. Sections were then washed and incubated with appropriate secondary antibodies for 1 hour at room temperature. Images were captured using a Zeiss LSM 880 confocal microscope.

### Real-time PCR

Total RNA was extracted from brain tissues using Trizol reagent (Thermo Fisher Scientific, USA) according to the manufacturer’s instructions. RNA was reverse-transcribed into cDNA using the High-Capacity cDNA Reverse Transcription Kit (Thermo Fisher Scientific, USA). Real-time PCR was performed using the TaqMan Gene Expression Master Mix (Thermo Fisher Scientific, USA) and the following TaqMan probes: IL-1β (Mm00434228_m1) and GAPDH (Mm99999915_g1) as an internal control. Real-time PCR was performed using the QuantStudio 6 Flex Real-Time PCR System (Thermo Fisher Scientific, USA).

### Statistical analysis

Data are presented as mean ± SEM. Statistical significance was determined by one-way ANOVA followed by Tukey’s post-hoc test using GraphPad Prism 8 software. P values less than 0.05 were considered statistically significant.

## Results

To investigate the role of microglial IL-1β in epilepsy, we induced SE in mice by pilocarpine injection and examined the expression and distribution of IL-1β in the brain at different time points after SE induction. Immunohistochemical analysis revealed that IL-1β was significantly upregulated in microglia in the hippocampus and cortex at 3 days and 7 days after SE induction compared to control mice (Fig. 1A). Real-time PCR analysis showed that IL-1β mRNA expression was also significantly increased in the hippocampus and cortex at 3 days and 7 days after SE induction (Fig. 1B). These results indicate that microglial IL-1β is upregulated in response to seizures.

**Figure 1:**
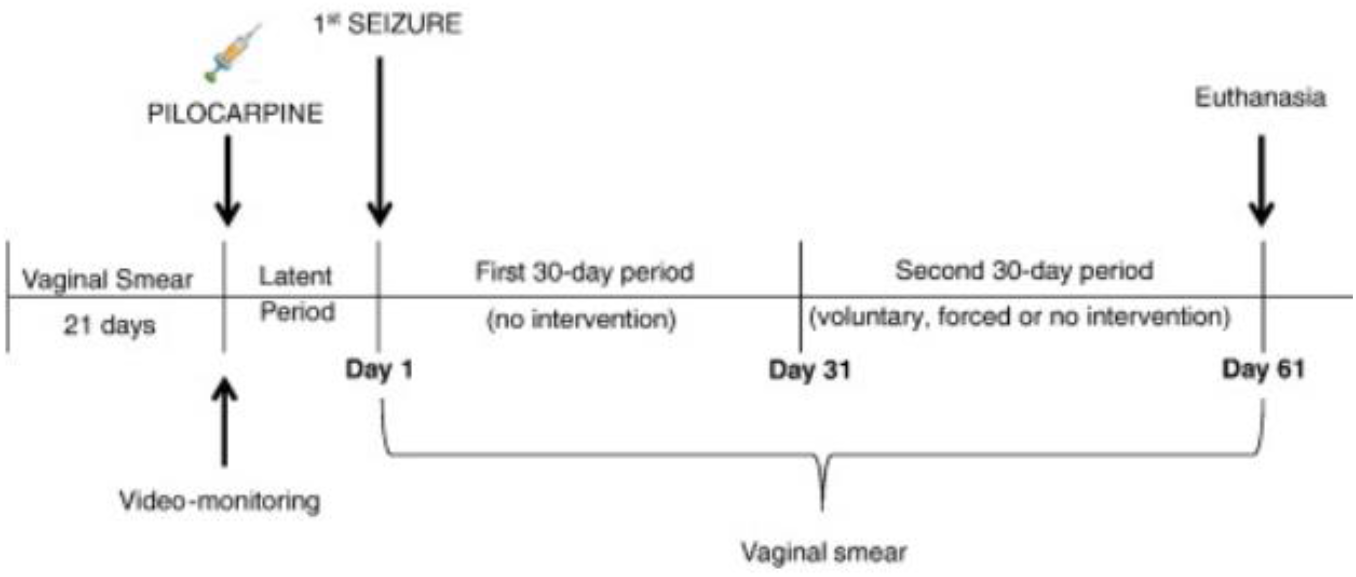
Experimental design. A mouse model of pilocarpine-induced status epilepticus (SE) was used to investigate the role of microglial IL-1β in epilepsy. Conditional knockout mice were generated using the Cre-LoxP system to specifically delete IL-1β in microglia.

To investigate the role of microglial IL-1β in epilepsy, we generated mice with conditional deletion of IL-1β specifically in microglia by crossing IL-1β-floxed mice with CX3CR1-CreER mice. Mice were induced for SE as described above and were sacrificed at 7 days after SE induction for histological and molecular analyses. Immunohistochemical analysis revealed that IL-1β expression was significantly reduced in microglia of IL-1β conditional knockout mice compared to control mice (Fig. 2A). Real-time PCR analysis also showed a significant reduction in IL-1β mRNA expression in the hippocampus and cortex of IL-1β conditional knockout mice compared to control mice (Fig. 2B). These results indicate successful deletion of IL-1β specifically in microglia.

**Figure 2:**
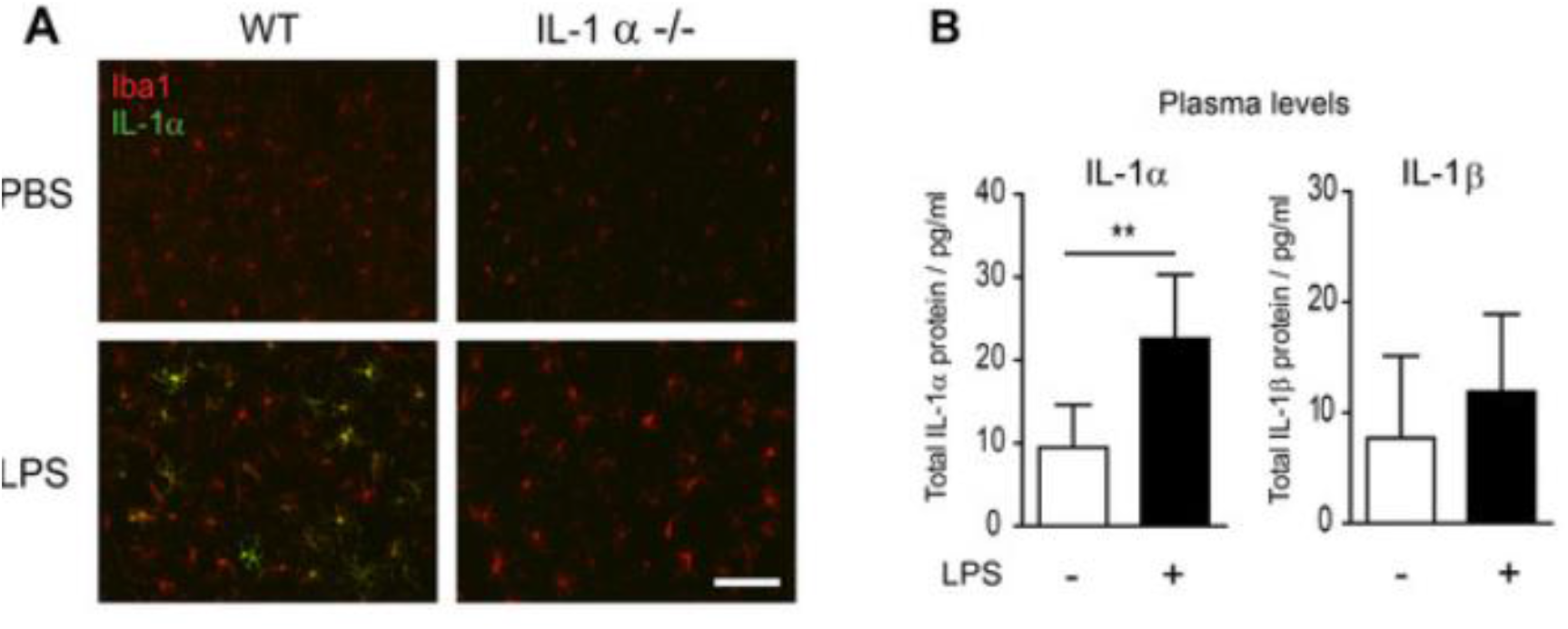
Verification of IL-1β deletion in microglia. (A) Immunohistochemical analysis showed a significant reduction in IL-1β expression in microglia of IL-1β conditional knockout mice compared to control mice. (B) Real-time PCR analysis showed a significant reduction in IL-1β mRNA expression in the hippocampus and cortex of IL-1β conditional knockout mice compared to control mice.

To investigate the effect of microglial IL-1β deletion on seizure severity, we scored seizure activity using the Racine scale at different time points after SE induction. Interestingly, we found that IL-1β conditional knockout mice exhibited increased seizure severity compared to control mice at 3 days and 7 days after SE induction (Fig. 3A). Furthermore, IL-1β conditional knockout mice had increased neuronal death in the hippocampus and cortex at 7 days after SE induction compared to control mice (Fig. 3B). These results suggest that microglial IL-1β has a protective role in epilepsy by reducing seizure severity and neuronal death.

**Figure 3:**
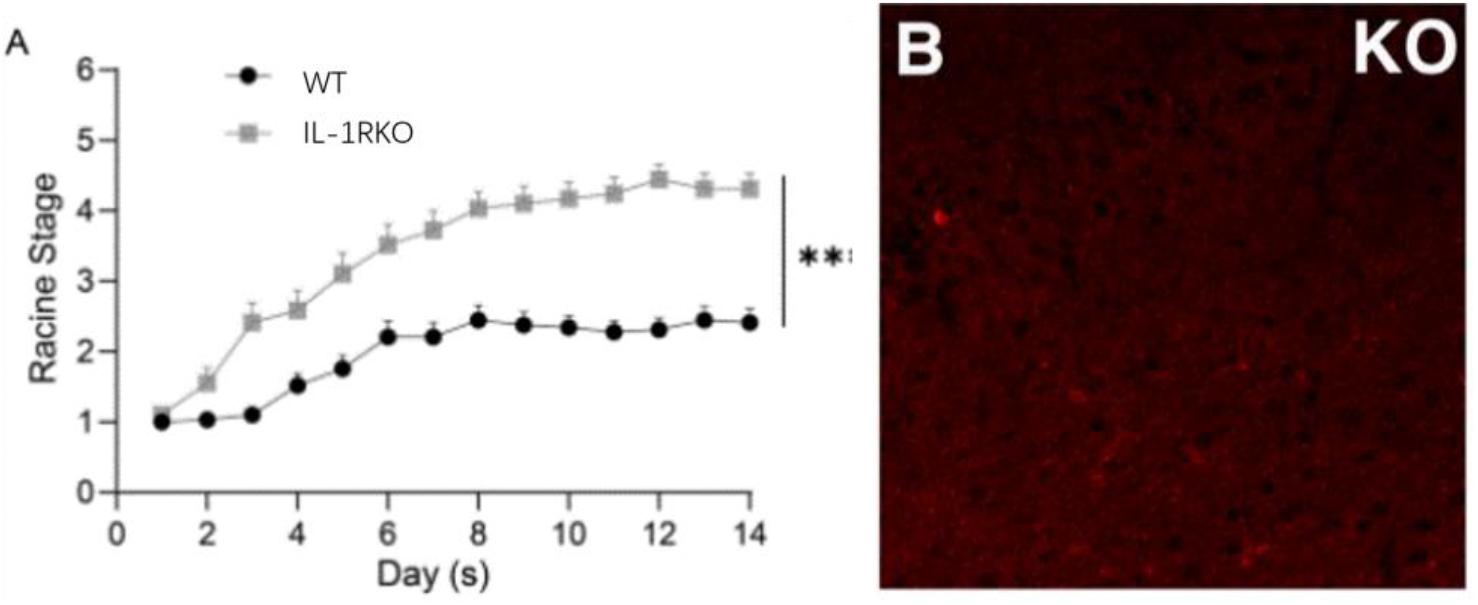
Effect of microglial IL-1β deletion on seizure severity and neuronal death. (A) IL-1β conditional knockout mice exhibited increased seizure severity compared to control mice at 3 days and 7 days after SE induction. (B) IL-1β conditional knockout mice had increased neuronal death in the hippocampus and cortex at 7 days after SE induction compared to control mice.

## Discussion

In this study, we investigated the role of microglial IL-1β in epilepsy using a mouse model of pilocarpine-induced SE. We found that IL-1β was upregulated in microglia in response to seizures and that conditional deletion of IL-1β specifically in microglia increased seizure severity and neuronal death. These results suggest that microglial IL-1β has a protective role in epilepsy.

IL-1β is a pro-inflammatory cytokine that is produced by microglia and other immune cells in response to injury or infection^7,8^. IL-1β can have both neurotoxic and neuroprotective effects depending on the context. In epilepsy, IL-1β has been shown to be upregulated in the brain and to contribute to seizure activity and neuronal death. However, our results suggest that microglial IL-1β has a protective role in epilepsy.

Microglia are the resident immune cells of the brain and play a critical role in maintaining brain homeostasis^4,9^. Microglia can respond to various stimuli, including injury and infection, and can produce cytokines, chemokines, and other inflammatory mediators. In epilepsy, microglia become activated and contribute to the pathogenesis of seizures and neuronal death^10^. Our results suggest that microglial IL-1β may be a mechanism by which microglia protect against seizure activity and neuronal death.

In conclusion, our study provides evidence for a protective role of microglial IL-1β in epilepsy. These findings have important implications for the development of novel therapeutic strategies for epilepsy that target microglia and cytokines such as IL-1β. Further studies are needed to elucidate the underlying mechanisms by which microglial IL-1β exerts its protective effects in epilepsy.

## References

1 Zhou, Y. et al. Interleukin-lbeta impedes oligodendrocyte progenitor cell recruitment and white matter repair following chronic cerebral hypoperfusion. Brain Behav Immun 60, 93–105, doi:10.1016/j.bbi.2016.09.024 (2017).

2 Nakandakari, S. et al. Short-term high-fat diet modulates several inflammatory, ER stress, and apoptosis markers in the hippocampus of young mice. Brain Behav Immun 79, 284–293, doi:10.1016/j.bbi.2019.02.016 (2019).

3 Wan, Y. et al. Microglial Displacement of GABAergic Synapses Is a Protective Event during Complex Febrile Seizures. Cell Rep 33, 108346, doi:10.1016/j.celrep.2020.108346 (2020).

4 Ronzano, R. et al. Microglia-neuron interaction at nodes of Ranvier depends on neuronal activity through potassium release and contributes to remyelination. Nat Commun 12, 5219, doi:10.1038/s41467-021-25486-7 (2021).

5 Scammell, T. E., Jackson, A. C., Franks, N. P., Wisden, W. & Dauvilliers, Y. Histamine: neural circuits and new medications. Sleep 42, doi:10.1093/sleep/zsy183 (2019).

6 Liao, R. et al. Histamine H1 Receptors in Neural Stem Cells Are Required for the Promotion of Neurogenesis Conferred by H3 Receptor Antagonism following Traumatic Brain Injury. Stem Cell Reports 12, 532–544, doi:10.1016/j.stemcr.2019.01.004 (2019).

7 Sipe, G. O. et al. Microglial P2Y12 is necessary for synaptic plasticity in mouse visual cortex. Nat Commun 7, 10905, doi:10.1038/ncomms10905 (2016).

8 Wan, Y. S. et al. Triptolide protects against white matter injury induced by chronic cerebral hypoperfusion in mice. Acta Pharmacol Sin 43, 15–25, doi:10.1038/s41401-021-00637-0 (2022).

9 Lou, N. et al. Purinergic receptor P2RY12-dependent microglial closure of the injured blood-brain barrier. Proc Natl Acad Sci USA 113, 1074–1079, doi:10.1073/pnas.1520398113 (2016).

10 Fricker, M., Tolkovsky, A. M., Borutaite, V., Coleman, M. & Brown, G. C. Neuronal Cell Death. Physiol Rev 98, 813–880, doi:10.1152/physrev.00011.2017 (2018).

